# Tegument ultrastructure in mother sporocysts of *Bunocotyle progenetica* (Markowski, 1936) Chabaud & Buttner, 1959 (Digenea: Hemiuridae)

**DOI:** 10.1101/2023.04.25.538318

**Authors:** I.M. Podvyaznaya, S.A. Filimonova

## Abstract

This investigation is devoted to the fine structure of mother sporocysts, the least studied stage of the digenean life cycle. The ultrastructure of the tegument of mature mother sporocysts of *Bunocotyle progenetica* was examined by transmission electron microscopy and described in detail. The tegument of these parthenitae is represented by an outer anucleate syncytium connected with the deeper-lying tegumental cells by cytoplasmic bridges. Its outer plasma membrane forms minute leaf-like outgrowths and numerous deep invaginations in the shape of interconnected channels. These channels, which pass in various directions and permeate almost the entire outer syncytium, considerably amplify its surface area. The cytoplasm of the outer layer of the tegument contains large mitochondria, microtubules and rare dense secretory granules, whose contents are discharged into the lumen of the channels. Numerous pinocytotic vesicles originate from the plasma membrane of the channels. Small endocytic vesicles are transported along the cytoplasmic bridges to the tegumental cells, where endocytosed food material is sorted and broken down. These cells are characterized by a well-developed Golgi apparatus, which is represented by multiple stacks of cisternae, and the presence of numerous endosomes at different stages of maturity and residual bodies. Some steps of endosomal maturation in the tegumental cells were traced. In addition to their digestive activity, tegumental cells produce secretory granules, which are transported to the outer syncytium after their maturation. It was shown for the first time that in mature parthenitae, the population of tegumental cells could be renewed at the expense of a reserve pool of undifferentiated cells. The ultrastructural features of the tegument of mother sporocysts of *B. progenetica* are discussed in the light of the concept of the enhanced trophic function of the tegument in sporocysts lacking the alimentary canal.

## INTRODUCTION

Mother sporocysts develop from miracidia larvae that have successfully infected the first intermediate host, the mollusc. Germinal cells formed in the mother sporocysts give rise to the individuals of the daughter parthenogenetic generation. Though the stage of the mother sporocyst is present in the life cycle of most digenean species (see reviews in Ginetsinskaya 1988; Galaktionov and Dobrovolkij 2003; Ataev 2017), it remains very poorly studied. For some species, its morpho-biological features have not been fully described. The descriptions of the mother sporocysts available in the literature are usually brief, fragmentary and based only on light microscopic observations. The few ultrastructural studies of the mother sporocysts are mostly devoted to the transformations of the tegument at the initial stages of their development in the host or *in vitro* (Southgate 1970; Basch and DiConza 1974; Køie et al. 1976; Køie and Frandsen 1976; Meuleman et al. 1978; Threadgold 1984; Matthews and Matthews 1991; Pan 1996; Poteaux et al. 2023).

In this study we described the fine structure of the tegument in mature, actively reproducing mother sporocysts of the hemiurid digenean *Bunocotyle progenetica* (Markowski, 1936) Chabaud & Buttner, 1959, a monoxenous parasite of the marine mud snails *Peringia ulvae* (Pennant, 1777) and *Ecrobia ventrosa* (Montagu, 1803). The mother as well as the daughter sporocysts of all digenean species completely lack the alimentary canal. This means that the uptake of nutrients by these parasites is possible only owing to the trophic activity of their tegument. In our study we attempted to trace the relationship between the functional features of the tegument and its fine organization in mother sporocysts of *B. progenetica*.

## MATERIAL AND METHODS

Mature sporocysts of *Bunocotyle progenetica* containing redial embryos in the brood cavity were obtained from mud snails *Peringia ulvae*. The snails were collected in summer (August) and during hydrological winter (March) in the Sukhaya Salma Inlet of the Chupa Bay of the White Sea. The ‘winter’ collection has been described in detail in Galaktionov and Podvyaznaya (2019).

The material for transmission electron microscopy was fixed with cold 3% glutaraldehyde on 0.1 М cacodylate buffer (pH=7.4) and subsequently (after 8-10 day) post-fixed with 1% osmium tetroxide solution on the same buffer at 4°C. Sucrose was added to the fixators and the wash buffer to obtain an osmolarity of 760 mOsm. The samples were then dehydrated and embedded into Epon-Araldite mixture following a standard procedure (Mollenhauer 1964). Thin and semi-thin sections of the sporocysts were made with the help of Leica EM UC6rt ultratome. Thin sections were stained with an aqueous solution of uranyl acetate and lead citrate and examined under Morgagni 268 transmission electron microscope operating at 80 kV. Besides, living sporocysts were examined with the help of Leica *DMLS* light microscope.

## RESULTS

Mature mother sporocysts of *Bunocotyle progenetica* are fairly motile worm-like organisms (1000-1700 µm long) residing in the hemocoel of the molluscan host. They have a well-developed brood cavity and a thin body wall comprising the tegument (Fig. 1A). The latter consists of the outer cytoplasmic layer and deeper-lying tegumental cells connected with the outer anucleate syncytium by cytoplasmic bridges (Fig. 1A). The covers of the sporocysts collected in summer and in winter did not have any noticeable ultrastructural differences.

**Fig. 1.**
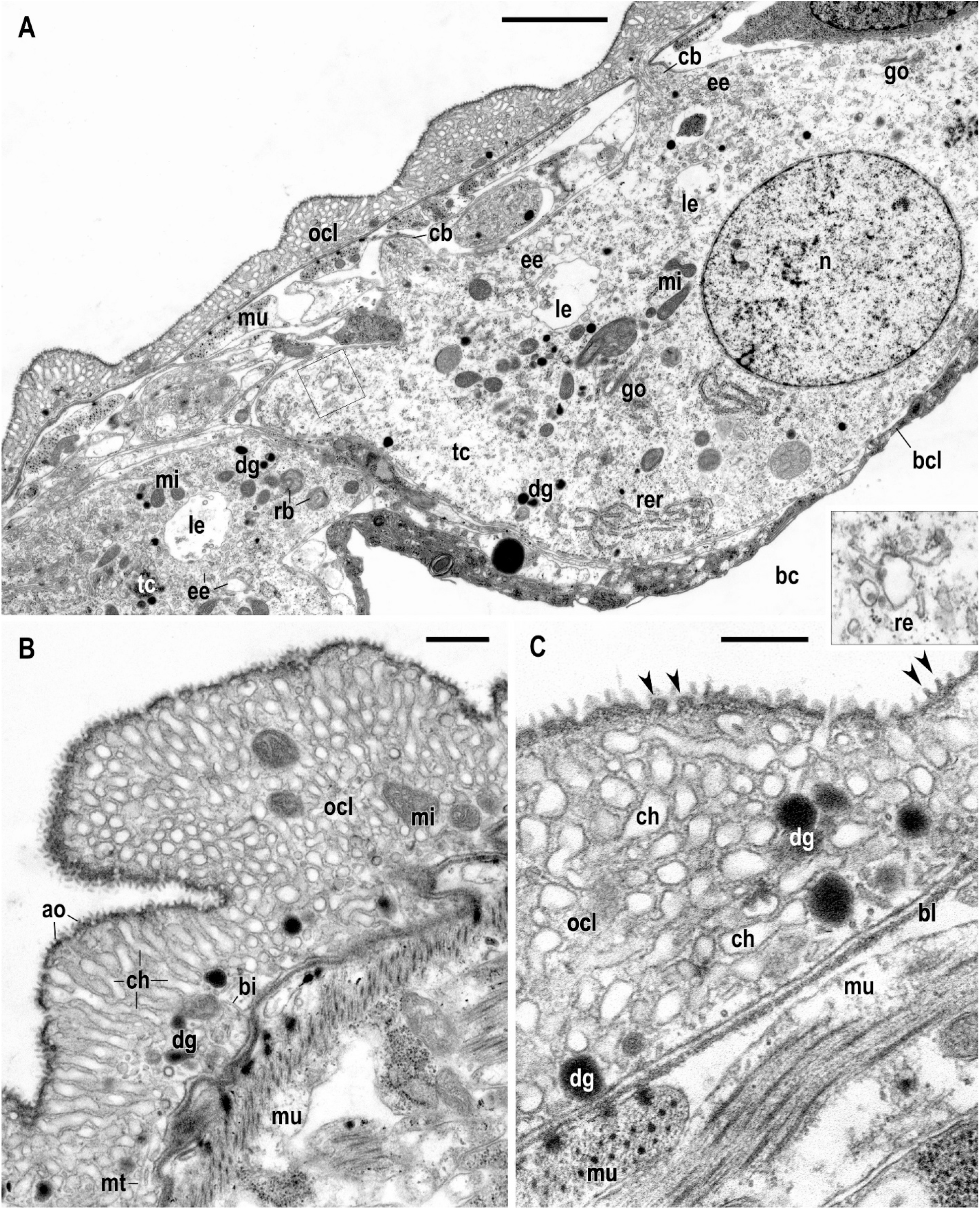
Tegument of mother sporocysts of *Bunocotyle progenetica*. A. General view of the sporocyst body wall with tegumental cells and outer cytoplasmic layer (OCL). Insertion shows an enlarged portion of the cytoplasm of a tegumental cell with a putative recycling endosome. B, C. Ultrastructural details of OCL; arrows point to different planes of sections through apical outgrowths. Abbreviations: ao – apical outgrowths; bc – brood cavity; bcl – brood cavity lining; bi – basal invaginations; bl - basal lamina; cb – cytoplasmic bridge; ch – channels; dg – dense granules; ee – early endosomes; go – Golgi apparatus; le – late endosomes; mi – mitochondria; mu – muscles; n – nucleus; ocl – OCL; rb – residual bodies; re – recycling endosome; rer – rough endoplasmic reticulum; tc – tegumental cell. Scale bars: A = 2 µm; B = 200 nm; C = 300 nm.

**Outer cytoplasmic layer** (OCL) apically bears dense minute leaf-like outgrowths (Fig. 1B, C). The cytoplasm within these outgrowths and a thin cytoplasmic layer just under them have an increased electron density owing to the concentration of fine electron-dense microfibrils. Besides the outgrowths, the outer plasma membrane of OCL forms numerous deep invaginations (Fig. 1B, C). They are represented by a network of interconnected channels passing in various directions and permeating almost the entire syncytium. These channels, about 120 nm in diameter, are unevenly filled with a substance of moderate electron density (Fig. 2A). Coated pinocytotic vesicles are formed from numerous coated pits in the plasma membrane limiting the channels (Fig. 2A, B). These vesicles, about 80 nm in diameter, are mainly located in the immediate vicinity of the channels’ membranes. The density of their contents is similar to that of the material filling the channels’ lumen (Fig. 2A, B). An accumulation of spherical or slightly irregular vesicles with a smooth limiting membrane is observed in the basal part of OCL; they are also about 80 nm in diameter (Fig. 2A). Numerous similar, though sometimes larger vesicles are present in the cytoplasmic bridges and in the adjacent areas of cytoplasm of the tegumental cells (Fig. 1A, 2B). In the present work, we regard them as endocytic vesicles.

**Fig. 2.**
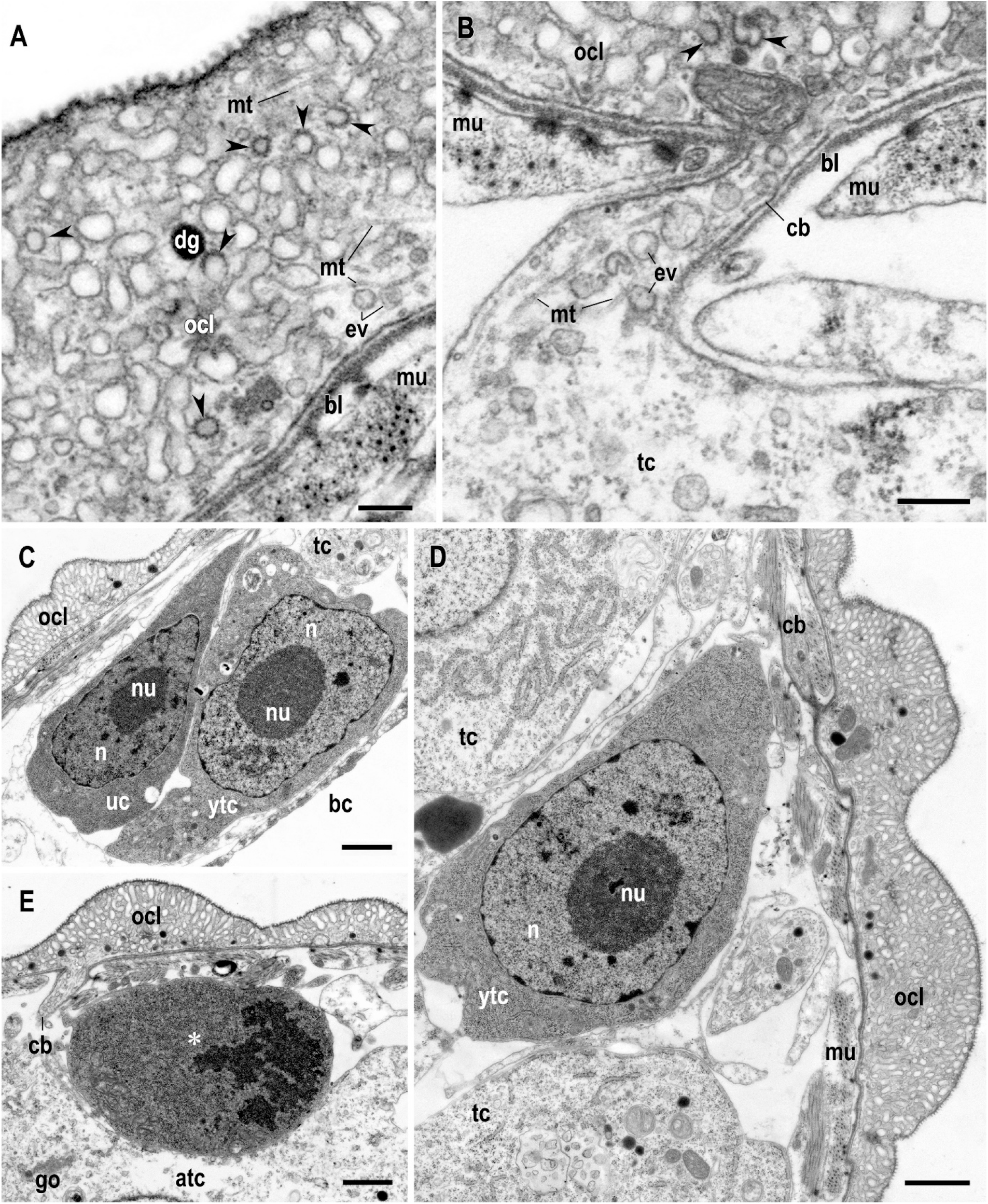
Pinocytosis in outer cytoplasmic layer (OCL) and renewal of population of tegumental cells in mother sporocysts of *Bunocotyle progenetica*. A. Enlarged portion of OCL showing formation of pinocytotic vesicles from coated pits (arrowheads) and endocytic vesicles in the basal part of OCL. B. Details of the cytoplasmic bridge. C. Undifferentiated and young tegumental cells. D. Early stage of differentiation of the tegumental cell. E. Mitosis (asterisk) of the undifferentiated cell. Abbreviations: atc – ageing tegumental cell; bc – brood cavity; bl - basal lamina; cb – cytoplasmic bridge; ch – channels; dg – dense granules; ee – early endosomes; ev – endocytic vesicles; go – Golgi apparatus; mi – mitochondria; mt – microtubules; mu – muscles; n – nucleus; nu – nucleolus; ocl – OCL; rb – residual bodies; rer – rough endoplasmic reticulum; ytc – young tegumental cell. Scale bars: A = 200 nm; B = 300 nm; C, D = 1 µm; E = 500 nm.

Secretory granules about 150 nm in diameter, bounded by a membrane and containing electron-dense finely fibrillar material (hereafter, dense granules), are found in OCL in much smaller numbers (Fig. 1B, 2D, 3A). These granules were regularly observed in direct contact with the membrane of the inner channels, and the lumen of the channels often contained electron-dense material similar to their contents (Fig. 3A).

**Fig. 3.**
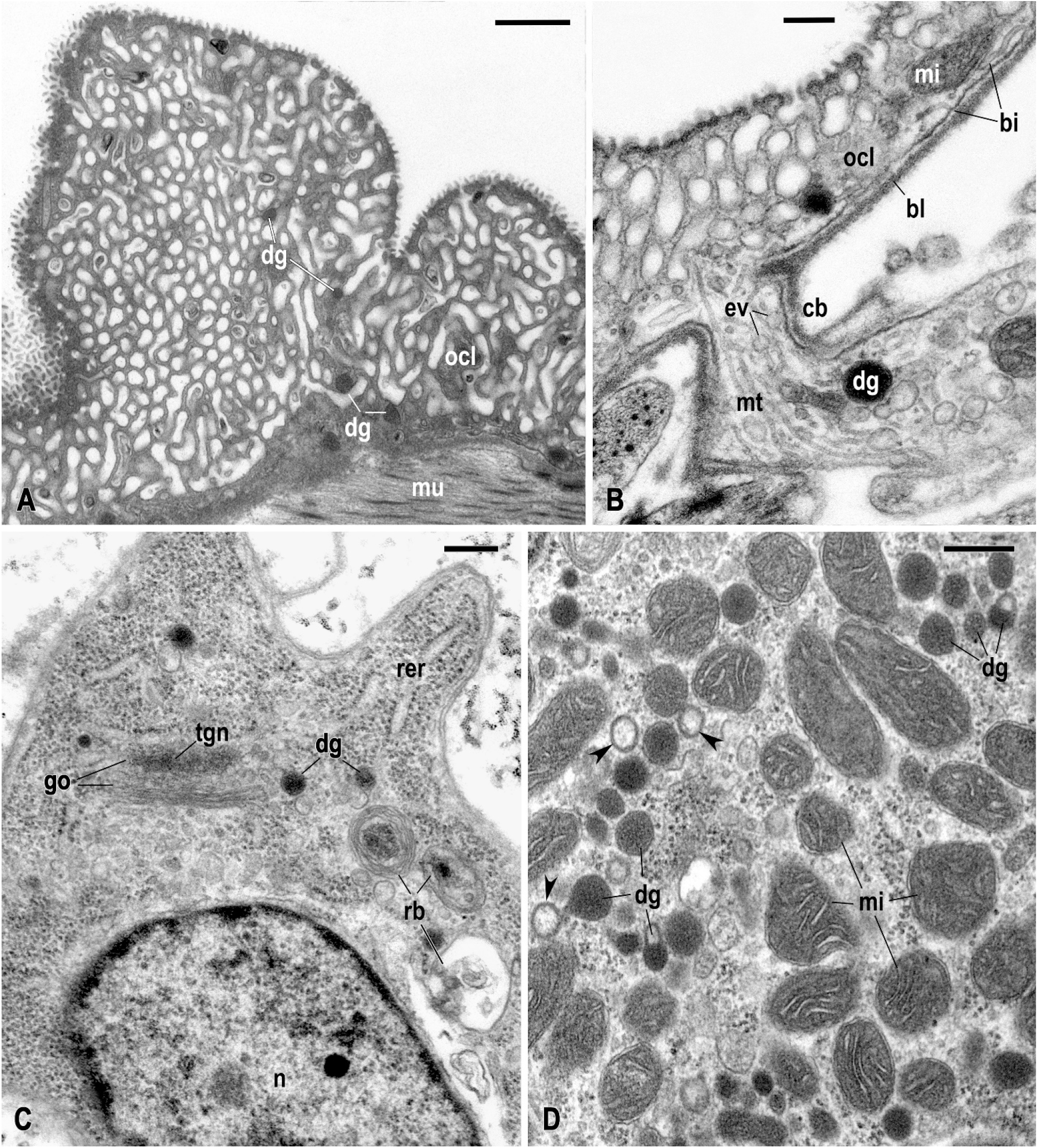
Dense granules and secretory process in the tegument of mother sporocysts of *Bunocotyle progenetica*. A. Release of secretory material (arrowheads) into the channels of outer cytoplasmic layer (OCL). Dense granules ready to discharge their contents are marked. B. Dense granule within the cytoplasmic bridge. C. Formation of secretory granules in a young tegumental cell. D. Maturation of secretory granules in the cytoplasm of the tegumental cell. Arrows point to “light” vesicles detached from the maturing granules. Abbreviations: bi – basal invaginations; bl - basal lamina; cb – cytoplasmic bridge; ch – channels; dg – dense granules; go – Golgi apparatus; mi – mitochondria; mt – microtubules; n – nucleus; ocl – OCL; rb – residual bodies; rer – rough endoplasmic reticulum; tgn – trans Golgi Network. Scale bars: A = 500 nm; B – D = 200 nm.

There are few organelles in the narrow interlayers of the cytoplasm between the inner channels of OCL. Apart from the vesicles and the inclusions described above, they contained only microtubules and rare large mitochondria, often located in the basal part of the outer syncytium (Fig. 1B, 2A). The basal plasma membrane of OCL forms sparse lamellar invaginations into the cytoplasm. It is underlain by a well-defined basal lamina, which also extends to the surface of the cytoplasmic bridges (Fig. 1B, C, 2A, B, 3B).

**Cytoplasmic bridges** connecting OCL and tegumental cells are numerous in mature sporocysts and have a relatively constant fine structure. It is characterized by the accumulation of long microtubules oriented along the axis of the bridge; vesicles, most of which are similar to the endocytic vesicles of OCL, are distributed between these microtubules (Fig. 2B, 3B). Occasionally, dense granules and mitochondria could be seen inside the bridges.

**Tegumental cells** are the most numerous cellular elements of the sporocyst body wall (Fig. 1A, 2C-E). Their ultrastructure changes noticeably with age. In particular, the age of a cell determines its size, the degree of development of the Golgi apparatus (GA) and rough endoplasmic reticulum (RER), the number of free ribosomes, dense granules and various vesicles.

In all specimens of mother sporocysts examined in our study, rare mitotic figures and a few small undifferentiated cells and tegumental cells at the early stages of specialization were found in the body wall near OCL (Fig. 2C-E). Tegumental cells at the early stages of specialization (Fig. 2D) differ from undifferentiated cells in having a larger nucleus with a larger nucleolus; besides, some RER and GA profiles and, occasionally, dense granules can be seen in their cytoplasm, which is densely filled with free ribosomes (Fig. 2C, D). These cells often have thin outgrowths, which are directed towards OCL but do not yet form cytoplasmic syncytial connections with it.

Young active tegumental cells are rather large and irregular in shape (Fig. 3C, 4A); each cell is connected with the OCL by several cytoplasmic bridges. The cells located far from the brood cavity (in the anterior body region) may also have outgrowths directed towards the cavity lining but not forming cytoplasmic bridges. The nuclei of young tegumental cells usually contain a small amount of condensed chromatin, which forms a thin layer under the nuclear envelope, one or two large nucleoli and numerous small RNP particles, which are distributed within the nucleoplasm (Fig. 2C, 3C, 4A). The number of mitochondria and RER profiles in the cytoplasm increases in these cells, but the concentration of free ribosomes is still high (Fig. 3C, 4A). Owing to this feature, young tegumental cells stand out as being denser than the others. GA is the best-developed organelle in their cytoplasm (Fig. 3C, 4A, B). It is represented by multiple large stacks of very narrow cisternae surrounded by an accumulation of diverse vesicles (Fig. 3C, 4B). Small transparent vesicles and vacuoles are often found close to the *cis* face of GA. A developed network of tubules tightly adjoining each other and passing in various directions is located at the *trans* side of GA (Fig. 3C, 4B). The tubular elements of this trans-Golgi network (TGN) are usually filled with material of an increased electron density. Most of the Golgi-derived vesicles are not more than 50 nm in size, and their contents are of a moderate electron density (Fig. 4B). Similar vesicles are often observed in contact with multivesicular bodies (MVBs) (Fig. 4C). Dense granules of secretory nature are produced by GA in smaller numbers. These granules often occur in accumulations in the cytoplasm of young cells (Fig. 3D, 4A). Immature secretory granules may be spherical, elongated, dumbbell-shaped or drop-shaped (Fig. 3D). As they mature, vacuoles with lighter contents are detached from them. These detached vacuoles have a narrow dense layer lining their membrane from the inside (Fig. 3D). Mature dense granules observed within the cytoplasmic bridges and OCL are usually spherical and homogeneous (Fig. 1C, 2A, D, 3A, B), though sometimes they may have a small light halo.

**Fig. 4.**
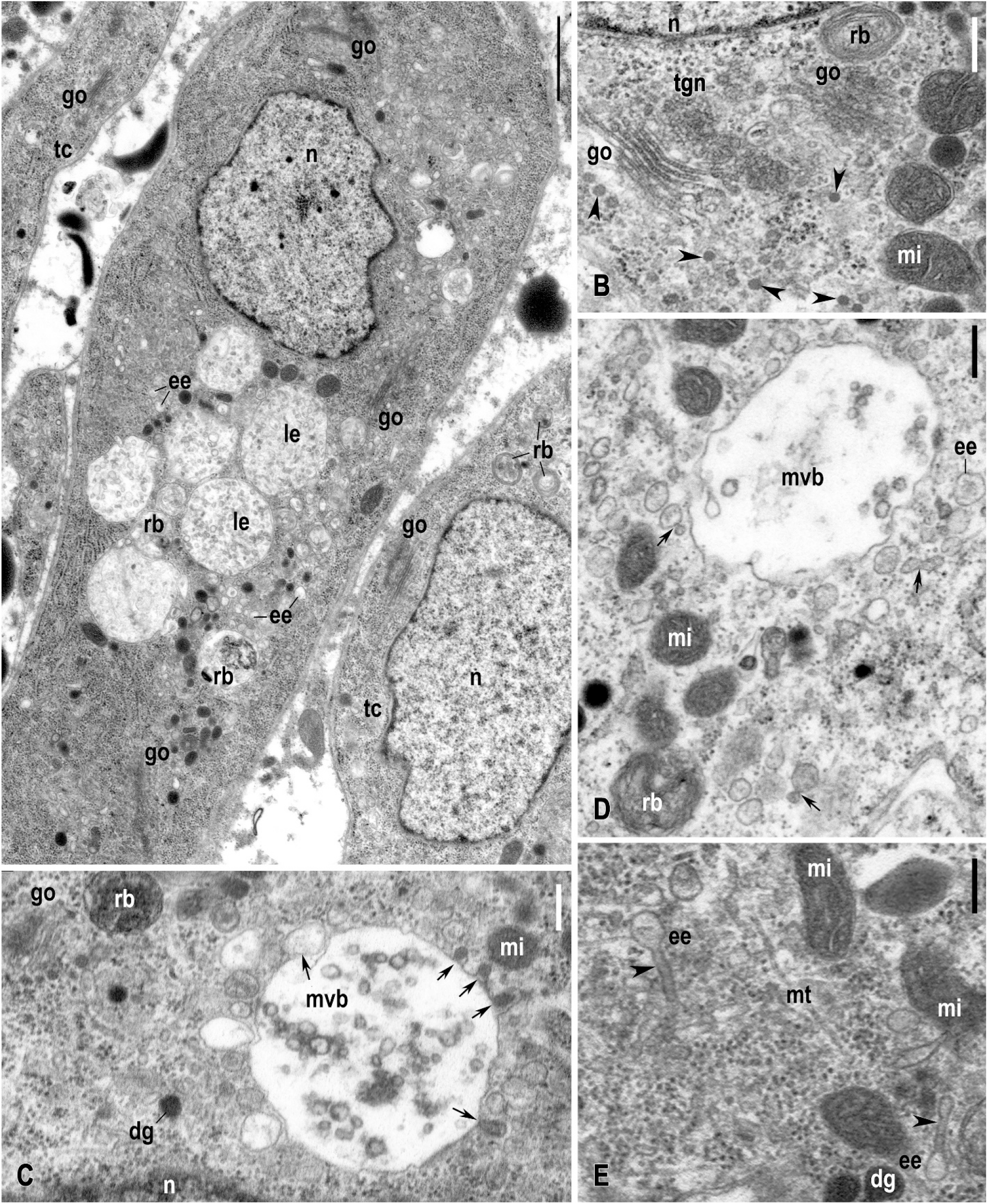
Maturation of endosomes in tegumental cells of mother sporocysts of *Bunocotyle progenetica*. A. General view of an active young tegumental cell with numerous endosomes and residual bodies. B. Golgi apparatus and small Golgi-derived vesicles with contents of moderate electron density (arrowheads). C, D. Multivesicular bodies (MVBs), small early endosomes and Golgi-derived vesicles in cytoplasm of the tegumental cells. Arrows point to pictures of presumable fusion of various vesicular structures. E. Early endosomes with tubular extensions. Abbreviations: dg – dense granules; ee – early endosomes; go – Golgi apparatus; mi – mitochondria; mt – microtubules; mvb – MVBs; n – nucleus; rb – residual bodies; tgn – trans Golgi Network; tc – tegumental cell. Scale bars: A = 1 µm; B, C, E = 200 nm; D = 300 nm.

Ageing tegumental cells are characterized by a light nucleus with a weakly pronounced nucleolus (Fig. 1A); the nuclei of some old cells show clear signs of degeneration. Their cytoplasm is considerably less dense (Fig. 2D, E) owing to a decreased concentration of free ribosomes and the profiles of RER and mitochondria. At the same time, their cytoplasm still contains multiple GA profiles. Similarly to the preceding cell cycle stages, the cytoplasm of these cells always contains microtubules and diverse vesicles and vacuoles, mostly represented by endosomes of various degree of maturity.

Endosomes in the cytoplasm of the tegumental cells are morphologically heterogeneous. Three main categories can be distinguished: early, recycling and late endosomes. Small early endosomes are especially numerous (Fig. 1A, 2B, 4D). Most of them are about 120 nm in diameter and have slightly uneven outlines in sections, resembling smaller vesicles observed in the cytoplasmic bridges. In the cytoplasm of the tegumental cells, small early endosomes can often be seen in close contact with each other and with larger vacuoles and vesicles (Fig. 4C, D). These pictures can be interpreted as the merging of maturing endosomes. Early endosomes also form tubular extensions (Fig. 4E), which make them look a bit like recycling endosomes. The latter are characterized by a stellar or an angular shape on sections and by more pronounced tubular elements (Fig. 1A, insertion), which can also be observed in the cytoplasm as separate structures. Late endosomes are represented by a population of MVBs, containing small intraluminal vesicles (ILVs) in the electron-light contents (Fig. 1A, 4A, C, D). Their size varies considerably, from 1 to 1.5 µm. Similarly to early endosomes, MVBs are often seen in close contact with vesicles of a smaller diameter, including endosomes at the previous stages of maturation; their fusion with small Golgi-derived vesicles has also been observed (Fig. 4C). Residual bodies in the cytoplasm of tegumental cells sometimes look like myelin-like bodies (or multilamellar bodies), while in other cases their composition and electron density is more heterogeneous (Fig. 1A, 3C, 4A-D). Many of the tegumental cells contain some autolysosomes with fragments of organelles inside (Fig. 1A).

## DISCUSSION

Our study showed that the general structural plan of the tegument of mother sporocysts of *Bunocotyle progenetica* is typical of digeneans. At the same time, some of its fine-structural features have never been recorded in studies of mother and daughter sporocysts of other digenean species.

We have already noted that in all generations of the sporocysts the uptake of the nutrients proceeds exclusively via the covers. As shown in previous ultrastructural studies (James et al. 1966; Southgate 1970; Køie 1971; Gibson 1974; Smith and Chernin 1974; Popiel 1978a,b; Matthews 1980; Fournier and Théron 1985; Žd’árská and Soboleva 1985; Al-Salman and James 1989; Pojmanska and Machaj 1991; Pan 1996; Russell-Pinto et al. 1996; Klag et al. 1997; Pinheiro et al. 2011; Podvyaznaya and Galaktionov 2012; Denisova et al. 2023), an intensification of the trophic function of the tegument in sporocysts is almost always accompanied by a considerable amplification of the area of their outer surface. In the majority of the digenean species examined in this respect, an increase in the absorptive surface area is achieved by means of numerous well-developed apical outgrowths of the outer plasma membrane, which may have a different shape. In mother sporocysts of *B. progenetica*, the area of the outer plasma membrane is increased manifold by means of a developed network of deep invaginations occupying a large part of the tegument volume, while small leaf-like apical outgrowths ensure only a marginal increase. These outgrowths are associated with a pronounced submembrane network of microfibrils, which appears to afford mechanical stability to the apical surface of the tegument. It cannot be ruled out that, besides being involved in absorption, the apical micro-outgrowths also have some mechanical function. For instance, they may perform a coarse filtration of the food material that gets into the inner channels of the tegument or prevent sliding during movement of the parasites along the small lacunae of the hemocoel.

An essentially similar ultrastructural organization of OCL has previously been described only once: in rediae of *Proterometra macrostoma* (see Uglem and Lee 1985). It is noteworthy that mature rediae of this species, similarly to the digenean sporocysts, lack the alimentary canal (Uglem and Lee 1985). The differences in the structure of OCL in parthenitae of *B. progenetica* and *P. macrostoma* are insignificant, being mostly related to the shape of the apical micro-outgrowths (finger-shaped in *P. macrostoma* and leaf-like in *B. progenetica*) and the arrangement of the inner channels (perpendicular to the basal surface of OCL in *P. macrostoma* and passing in various directions in *B. progenetica*). Uglem and Lee (1985) have estimated that the inner channels and the outer micro-outgrowths taken together amplify the surface area of the outer plasma membrane of the tegument of *P. macrostoma* more than 15-fold. They have also demonstrated experimentally (Uglem 1980; Uglem and Lee 1985) that in rediae of *P. macrostoma* glucose is actively transported across the plasma membrane limiting the inner channels. Literature data indicate (see review in Smyth and Halton 1983; Galaktionov and Dobrovolkij 2003) that the absorption of low molecular weight carbohydrates by the tegument with the help of active transport mechanisms as well as by simple or facilitated diffusion is also characteristic of sporocysts of some digenean species. In the lacunar system of the molluscs, mother sporocysts of *B. progenetica* are in constant contact with the hemolymph, containing large quantities of dissolved organic substances with a low molecular weight. It is highly likely that the transmembrane transport of some of these substances occurs in their covers.

In this study we showed that in the tegument of mother sporocysts of *B. progenetica*, the putative transmembrane transport is supplemented by a fairly active vesicular transport, owing to which large organic molecules can get into the parasite’s body. Evidence of pinocytosis has been found in the tegument of sporocysts of many digeneans, including the parthenitae of *Cercaria vullegeardi, Cercaria buccini, Cercaria helvetica XII, Fasciola hepatica, Schistosoma mansoni*,

*Meiogymnophallus minutus, Prosorhynchoides borealis, Podocotyle* sp. and others (Southgate 1970; Køie 1971; Reader 1974; Popiel 1978b; Meuleman et al. 1978; Al-Salman and James 1989; Podvyaznaya and Galaktionov 2012; Denisova et al. 2023). In most of these species, pinocytotic vesicles are formed at the apical surface of OCL in the areas free from outgrowths. In mother sporocysts of *B. progenetica*, pinocytotic vesicles are the derivatives of deep invaginations of the outer plasma membrane. Considering the large volume of the channels (invaginations) of OCL and the abundance of endosome-like vesicles in the tegumental cells, it can be assumed that several types of vesicular transport coexist in these parthenitae. Receptor-mediated endocytosis is easily discernible in sections owing to a characteristic clathrin coating of its vesicles. Deeper in the cytoplasm the clathrin coating disintegrates quickly, and these vesicles become indistinguishable from those that may arise from caveolae. The identification of caveolin vesicles in *B. progenetica* is hampered by the presence of numerous inner channels passing in various directions, with which they can be easily confused in the sections. Therefore, additional studies employing special markers are necessary to prove or disprove the existence of caveolin endocytosis in these sporocysts.

In *B. progenetica*, endocytic vesicles move to the cytoplasmic bridges with numerous microtubules, along which they are transported to the tegumental cells. We managed to observe some stages of endocytic pathway in the cytoplasm of the tegumental cells. According to the modern concept of the vesicular transport (Miaczynska and Stenmark 2008; Huotari and Helenius 2011; Kornilova 2014; Tu et al. 2020), transformation of endosomes involves the fusion of early endosomes and endocytic vesicles into larger endosomal structures and the formation of tubular extensions reflecting the sorting of their contents and the recycling of the components of the outer plasma membrane. Conversion of early endosomes into late endosomes can be considered as the next stage of endosomal maturation. In *B. progenetica*, late endosomes are represented by numerous large MVBs, which are capable of fusing with small vesicles with contents of moderate electron density. These Golgi-derived vesicles presumably contain lysosomal enzymes and may be classified as primary lysosomes. The presence of transitional forms between large MVBs and dense telolysosomes with heterogeneous contents suggests that the latter are the final stage of intracellular processing of endocytosed material. Myelin-like structures (multilamellar bodies), which were also observed in the tegumental cells, are possibly the end product of autophagy. In *B. progenetica*, residual bodies are not transported to OCL, as they are in sporocysts of some other digenean species (see, e.g., Al-Salman and James 1989; Podvyaznaya and Galaktionov 2012), but accumulate in tegumental cells; sometimes they discharge their contents into the intercellular space.

To sum up, in the tegument of mother sporocysts of *B. progenetica*, the processes associated with its trophic activity are spatially separated. OCL is adapted to capture as broad a range of nutrients as possible, including some large molecules, which cannot get into the cell across the plasma membrane. If caveolin endocytosis also occurs in the tegument, it additionally enhances the uptake of nutrients by increasing the rate of their passage across the absorptive surface. Tegumental cells are responsible for sorting and breakdown of endocytosed food material; from these cells the nutrients are transported into other tissues of the sporocyst in the available form.

It should be noted that the distribution of functions in the covers of the sporocysts found in our study is characteristic not of all the digeneans in which endocytosis has been described. In some species the ultrastructural features of the OCL indicate the activity of intracellular digestion (Reader 1974; Al-Salman and James 1989; Podvyaznaya and Galaktionov 2012). For instance, the OCL of *C. helvetica* XII, *M. minutus* and *P. borealis* contains polymorphic vesicles with heterogeneous contents of non-secretory nature, as well as residual bodies (Reader 1974; Al-Salman and James 1989; Podvyaznaya and Galaktionov 2012). Judging by the results of the cytochemical analysis of *C. helvetica* XII, many of these structures demonstrate the activity of acid phosphatase (Reader 1974), suggesting the functioning of the lysosome system of the cell in this part of the tegument. Another evidence indicating the functioning of this system is the presence of GA in the OCL of *P. borealis* and *C. helvetica* XII (Reader 1974; Podvyaznaya and Galaktionov 2012).

Age-related changes in the tegumental cells have been recorded in sporocysts of various species (see, e.g., Al-Salman and James 1989; Podvyaznaya and Galaktionov 2012; and others). At the same time, data on the differentiation of new tegumental cells in mature parthenitae have been lacking. Our observations indicate that mature mother sporocysts of *B. progenetica* have a pronounced capacity to the renewal of the population of tegumental cells at the expense of the reserve pool of undifferentiated cells. This feature of their tegument may be associated with a short life span of the tegumental cells. An active involvement of these cells in intracellular digestion appears to accelerate their ageing and degradation. At the same time, the mother sporocysts of *B. progenetica* may live for a fairly long time by the standards of this life-cycle stage (up to six months), their life span depending on the season when the mollusc has been infected (Levakin 2008). These parthenitae can overwinter in the body of the host; during this time they retain the ability to move actively, and the process of development of redial embryos continues in their brood cavity (Podvyaznaya and Galaktionov 2021). Apparently, this would have been impossible without the continued trophic function of the tegument.

## ACKNOWLEDGEMENTS

We are very grateful to Natalia Lentsman for her help in preparing the English version of the manuscript. This study was carried out using equipment of the Core Facilities Centre “Taxon” at the Zoological Institute of the Russian Academy of Sciences, St. Petersburg, Russia (http://www.ckp-rf.ru/ckp/3038/?sphrase_id=8879024), and was supported by budget funding of the Russian Academy of Sciences (project numbers 122031100260-0 and1021051603202-7).

